# Modeled Oxidative Lesions Remodel the ABCB1 Linker: A Hypothesis for Transporter Regulation

**DOI:** 10.64898/2026.10.10.758164

**Authors:** Mansoureh Hoseini, Hossein Abbasi

**Author notes:** Corresponding author. Corresponding Author’s Information: Hossein Abbasi.

## Abstract

Multidrug resistance mediated by P-glycoprotein (P-gp; ABCB1) remains a major barrier to chemotherapy. Cold atmospheric plasma (CAP) has increased drug responsiveness in resistant cancer models, but the molecular link between plasma-derived oxidative chemistry and ABCB1 regulation remains unresolved. We investigated whether site-specific oxidative lesions remodel the intrinsically disordered linker connecting nucleotide-binding domain 1 to transmembrane domain 2. Native-ensemble exposure analysis identified contrasting oxidation candidates: dynamically exposed Met653 and persistently exposed Trp698. Met653 oxidation to methionine sulfoxide and Trp698 oxidation to kynurenine were examined individually and together using a balanced 2×2 molecular-dynamics design with a mechanically tethered linker construct. Analyses integrated global descriptors, residue-resolved mobility, backbone order, native-state occupancies, projected breadth, and conformational switching.

Trp698→KYN produced the dominant response: all seven global descriptors shifted toward restriction, accompanied by distributed mobility changes, compaction, and increased intramolecular hydrogen bonding. In the radius-of-gyration/end-to-end-distance projection, compact-state occupancy increased from 46.9% to 99.7%, switching decreased from 6.50 to 0.07 transitions per 10 ns, and projected breadth contracted. Met653→SME produced weaker, heterogeneous effects, while the double-modified linker remained restricted but showed greater projected exchange than Trp698→KYN alone.

In this model, Trp698→KYN generated a compact, low-exchange projected ensemble in the tethered ABCB1 linker. These findings identify oxidation-site-specific linker remodeling as a candidate molecular intermediate that could influence linker-mediated domain coordination, while requiring experimental validation of lesion formation, transporter activity, and multidrug resistance outcomes.

## 1. Introduction

Multidrug resistance (MDR) remains a major cause of chemotherapy failure. Among its clinically important mechanisms, overexpression of P-glycoprotein (P-gp; ABCB1) lowers intracellular drug accumulation by exporting structurally diverse chemotherapeutic agents ^[1–5]^. Despite extensive efforts, direct P-gp inhibition has achieved limited clinical success because of insufficient selectivity, dose-limiting toxicity, unfavorable pharmacokinetic interactions, and compensatory resistance mechanisms ^[6–10]^. Alternative approaches capable of modulating transporter function without relying exclusively on competitive inhibition therefore remain of considerable interest.

Cold atmospheric-pressure plasma (CAP) has emerged as a potential adjunct to cancer therapy. CAP generates a complex physicochemical environment containing reactive oxygen and nitrogen species (RONS), charged particles, electric fields, and ultraviolet radiation ^[11–14]^. These components may act directly on cells or generate longer-lived reactive products in surrounding liquids. CAP and plasma-activated media have increased chemotherapy responsiveness in several resistant cancer models ^[15–19]^, but the underlying mechanisms are likely multifactorial and remain incompletely resolved.

ABCB1 is a plausible component of this response because its activity depends on coordinated conformational dynamics as well as expression and membrane localization. The transporter contains two transmembrane domains (TMDs), which form the substrate-binding and translocation pathway, and two cytosolic nucleotide-binding domains (NBDs), which bind and hydrolyze ATP ^[20–25]^. Communication between these domains drives transitions among inward-facing, nucleotide-bound, and outward-facing conformations. Perturbations that alter the timing, range, or coupling of these motions could consequently affect drug efflux without directly occupying the substrate- or ATP-binding sites.

The intrinsically disordered linker connecting NBD1 to TMD2 is an understudied component of this coupling network. Linker deletion, truncation, and sequence-replacement experiments have shown that its integrity and flexibility influence ATP hydrolysis, substrate transport, and communication between the two halves of P-gp ^[26,27]^. Molecular dynamics simulations further support a role for the linker in shaping nucleotide-binding domain arrangements and facilitating communication across the two halves of the transporter ^[28–31]^. Rather than functioning as a passive connector, the linker appears to support transport through a dynamic conformational ensemble. Its solvent exposure and rapidly interconverting transient contacts also make it a plausible sensor of chemical perturbation: a local change in side-chain polarity, geometry, or hydrogen-bonding capacity may redistribute compact, extended, and exchanging linker states ^[32–34]^.

Previous reduced-construct work showed that noncovalent interactions with selected CAP-associated small molecules can remodel the ABCB1 linker and that terminal constraints strongly shape whether these effects appear as compaction, altered conformational breadth, or reduced exchange [35,36]. These observations motivated the use of a mechanically tethered construct as a controlled model for testing persistent chemical perturbations under domain-mimetic terminal constraints.

Covalent oxidative lesions represent a distinct form of CAP-associated protein modification because their structural effects may persist after the initiating oxidant has disappeared. Methionine oxidation produces methionine sulfoxide, increasing side-chain polarity and hydrogen-bonding capacity, whereas tryptophan oxidation can generate kynurenine through disruption and opening of the indole ring [37,38]. These transformations may alter solvation, transient packing, and backbone preferences in context-dependent ways. Because multiple lesions may coexist, the ensemble produced by their combination cannot necessarily be inferred from their individual effects. To our knowledge, individual and combined oxidative modifications of the ABCB1 NBD1–TMD2 linker have not previously been examined at the conformational-ensemble level.

We selected two sites with contrasting solvent-exposure profiles in native reference simulations (Fig. 1). Met653 was variably exposed among trajectories, whereas Trp698 remained consistently accessible. Met653 oxidation was modeled as methionine sulfoxide (Met653→SME) and Trp698 oxidation as kynurenine (Trp698→KYN). Native, single-modified, and double-modified linkers were examined in a balanced 2×2 design.

**Figure 1.**
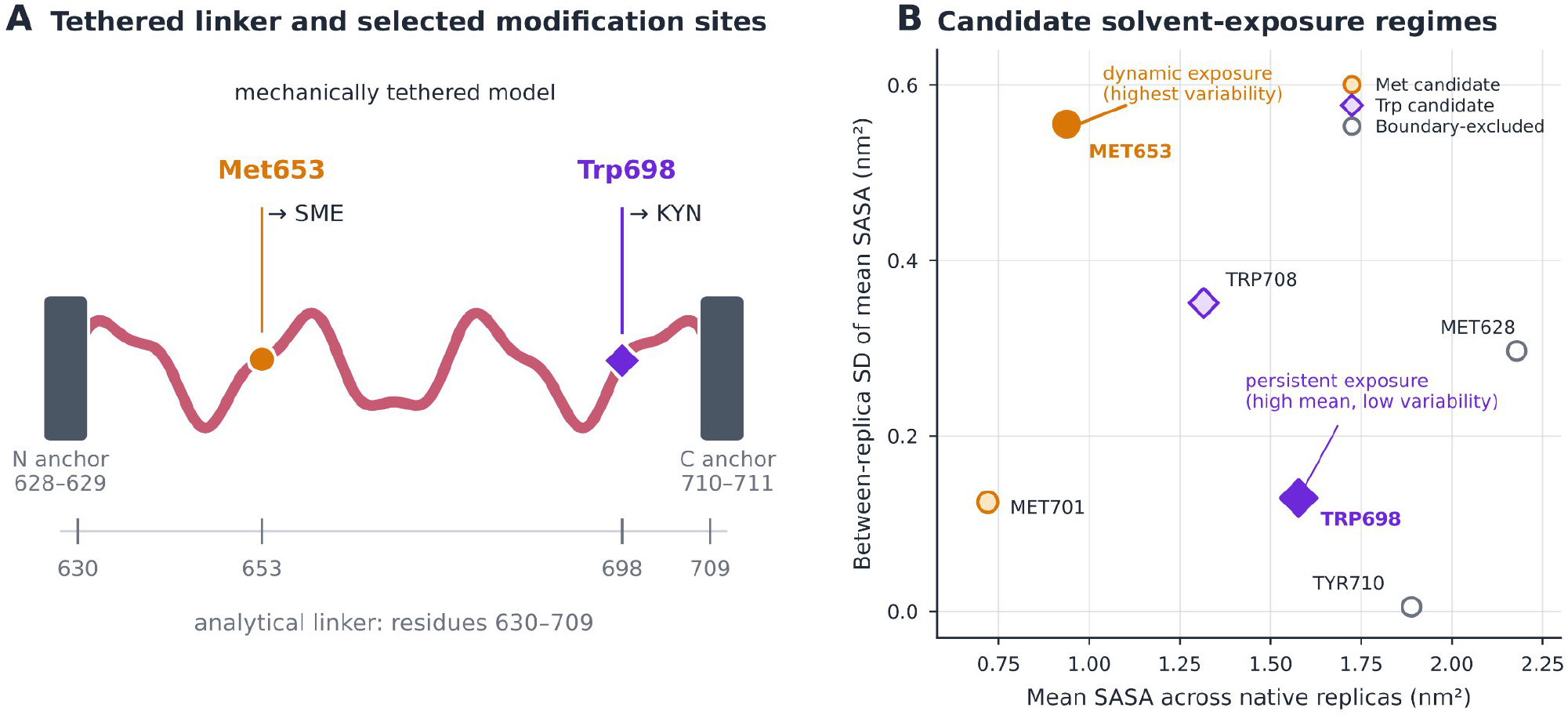
Selection of Met653 and Trp698 as contrasting oxidation sites in the tethered ABCB1 linker. (A) Sequence-positioned schematic of the mechanically tethered ABCB1 NBD1–TMD2 linker construct. Residues 630–709 constitute the analytical linker, whereas residues 628–629 and 710–711 form the excluded N- and C-terminal anchor segments. Met653 was modeled as methionine sulfoxide (SME), and Trp698 was modeled as kynurenine (KYN). (B) Mean whole-residue SASA plotted against the between-replica standard deviation of mean SASA across three native reference trajectories. Met653 represents dynamic exposure because it showed the greatest non-terminal between-replica SASA variability. Trp698 represents persistent exposure because it combined the highest non-terminal tryptophan SASA with comparatively low variability. Terminal residues are shown as boundary-excluded references.

We asked whether these defined modifications redistribute the mechanically constrained linker ensemble and whether the modeled modification states produce distinct responses. For outcomes with interpretable measurement scales, we also examined whether the combined modification deviated from raw-scale additivity. Analyses integrated global conformational descriptors with residue-resolved mobility and backbone order, projected landscapes, native-ensemble-defined state occupancies, conformational switching, and ensemble breadth. This reduced-construct study does not reproduce CAP treatment or measure full-length ABCB1 transport; it tests whether chemically defined oxidative lesions can generate persistent remodeling of the tethered linker ensemble.

## 2. Materials and Methods

### 2.1. ABCB1 linker construct and mechanical tethering

The tethered ABCB1 linker construct (residues 628–711; analytical linker Thr630–Pro709) and the four-restraint flat-bottom tether were used exactly as established in the preceding tethered-linker studies [35, 36]. Full restraint definitions are in Supplementary Method S1.

### 2.2. Selection of oxidation sites

Candidate sites were evaluated using residue-level solvent-accessible surface area (SASA) and RMSF data from three native tethered-linker reference trajectories over 100–200 ns. SASA was calculated with gmx sasa using the default 0.14-nm solvent probe; the reported value was the total SASA of the complete residue. Replica-mean SASA values were summarized by their mean and between-replica standard deviation. Six oxidation prone sulfur-containing or aromatic reference residues were considered: Met628, Met653, Met701, Trp698, Trp708, and Tyr710 (Fig. 1). Terminal Met628 and Tyr710 were excluded because they were outside the analytical linker. Met653 was selected as a dynamic-exposure site because it showed the largest between-replica variability among non-terminal candidates (mean SASA, 0.938 nm^2^; between-replica SD, 0.556 nm^2^). Trp698 was selected as a persistent-exposure site because it had the highest mean SASA among non-terminal tryptophans (1.577 nm^2^) with comparatively low variability (SD, 0.129 nm^2^). Trp708 was accessible but more variable and closer to the C-terminal boundary, whereas Met701 showed lower accessibility and variability. SASA was used solely to define contrasting exposure regimes; it was not interpreted as oxidation probability, reaction kinetics, or biological susceptibility.

Met653 and Trp698 were modeled as methionine sulfoxide and kynurenine, respectively [37,38]. These chemically established oxidative products were selected as defined representative lesions with contrasting physicochemical consequences and available CHARMM-GUI residue definitions [40]. The KYN residue represented L-kynurenine, whereas SME represented L-methionine-(R)-sulfoxide; the corresponding sulfur-S diastereomer was not examined. These modeled lesions were not assumed to be the exclusive or predominant products formed at these sites during CAP exposure.

### 2.3. Modified-residue models and factorial simulation design

Four tethered-linker conditions formed a balanced 2×2 design: native linker, Met653→SME, Trp698→KYN, and the double-modified linker. The SME and KYN residues were assigned using the corresponding CHARMM-GUI modified-residue definitions. All conditions were generated from the same initial tethered-linker conformation. For each condition, three independent production replicas were initiated from the same prepared coordinate model using independently generated velocities and random seeds.

### 2.4. System preparation and molecular-dynamics simulations

Systems were prepared with CHARMM-GUI at pH 7.4 (12.5 nm cubic box, CHARMM-modified TIP3P, ∼0.15 M KCl) retaining the CHARMM-GUI SME/KYN definitions [40]. Simulations used GROMACS 2024.2 with CHARMM36m [41, 42] and the equilibration protocol of [35] (Supplementary Method S1). Four modification states × three replicas produced 12 trajectories (2.4 μs); primary analyses used 100–200 ns.

### 2.5. Global conformational and mechanical descriptors

The seven prespecified global descriptors (Schlitter entropy, covariance trace, core Cα RMSF, effective spring constant, core S2min, persistence length, dihedral-transition count) and alignment procedures are identical to [35] (Supplementary Method S1).

Conformational entropy was estimated from the mass-weighted atomic covariance matrix using the Schlitter approximation and reported in J mol^−1^ K^−1^. The trace of the positional covariance matrix quantified the overall amplitude of collective fluctuations. C*α* RMSF was calculated for each residue and replica, and the mean across residues 630–709 was used as the replica-level global RMSF.

The effective spring constant of each trajectory was estimated using the equipartition theorem:

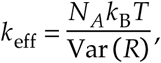

where *R* is the end-to-end distance and *T* = 310.15 K. This fluctuation-based stiffness applies within the imposed tethered boundary condition and is not an intrinsic material constant of the isolated sequence.

Backbone order parameters were calculated with gmx chi. For each residue, the GROMACS 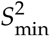 output was retained, and the mean across residues 630–709 was used as the replica-level global order descriptor; larger values indicate lower angular variability. Persistence length was obtained from backbone directional correlations and reported in backbone bonds. Backbone dihedral transitions were also quantified as a measure of local conformational exchange.

Radius of gyration, end-to-end distance, and intramolecular hydrogen-bond count were analyzed as supporting descriptors rather than components of the seven-metric primary analysis. Hydrogen bonds were defined by a donor–acceptor distance ≤ 0.35 nm and a hydrogen–donor–acceptor angular deviation ≤ 30^°^. Counts were averaged within each trajectory.

### 2.6. Residue-resolved remodeling analysis

Residue-resolved C*α* RMSF and backbone 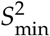 profiles were generated for residues 630–709. Each trajectory produced one 80-residue profile, and trajectories—not residues or frames—were treated as independent observations. 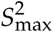 was retained only as a supporting descriptor because its values were close to saturation. Four contrasts were calculated for each residue:

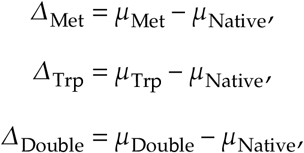

and

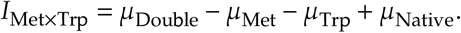

RMSF effects were oriented so that positive values denoted reduced mobility, whereas positive 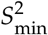 effects denoted increased backbone order. For uncertainty propagation, each condition mean was represented by a Student-*t* distribution with two degrees of freedom, centered on the sample mean and scaled by the replica standard error, 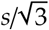. 3. 30,000 Monte Carlo draws propagated uncertainty through each contrast. Hedges *g* was calculated using the pooled within-condition replica standard deviation. Interpretation emphasized contiguous regions. A candidate RMSF or 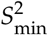 region required at least three consecutive residues for which every residue showed the same oriented direction, *g* ≥ 0.80, posterior probability of the observed direction ≥ 0.80, and at least 80% sign retention under balanced replica deletion. Two-metric consensus required at least two consecutive residues satisfying the candidate criterion for both RMSF and 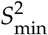 in the same restriction-oriented direction.

Two windows were specified before regional interpretation: residues 651–655 around Met653 and residues 696–700 around Trp698. Values were averaged within each window for every replica before inference. Welch–Satterthwaite *P* values and Benjamini–Hochberg-adjusted *q* values were calculated descriptively for individual residues. Because no isolated residue survived false-discovery-rate correction, individual residues were not interpreted as independently significant.

### 2.7. Projected conformational landscapes and state exchange

Conformational landscapes were projected onto the joint radius-of-gyration/end-to-end-distance coordinate, (*R*_*g*_ ,*R*). Each replica was represented on a common 28 × 28 grid. Replica histograms were normalized separately and averaged with equal weight, preventing trajectories or frames from being treated as independent replicates. The averaged probability surface was smoothed using a Gaussian kernel with a width of one grid bin. Projected free energy was calculated as

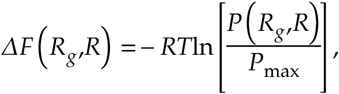

where *T* = 310.15 K and *RT* = 2.579 kJ mol^−1^. Adjacent 2 × 2 probability bins were aggregated for a 14 × 14 display grid; all quantitative calculations used the original 28 × 28 grid. The surfaces therefore represent two-coordinate projected potentials of mean force rather than complete macromolecular free-energy landscapes.

Compact, intermediate, and extended states were defined by *k*-means clustering with *k* = 3. Clustering used only standardized *R*_*g*_ and *R* values from the three native trajectories. Centroids were ordered by joint compactness and fixed before modified-condition frames were classified; state labels are therefore relative to the native projected ensemble. Every retained 10-ps frame was assigned to the nearest native centroid to calculate replica-level occupancies. For switching analysis, coordinates were averaged in 100-ps blocks, assigned to the nearest centroid, and passed through a three-block majority filter to suppress single-block boundary flicker. Transitions were reported per 10 ns. Distributional similarity to the native ensemble was quantified using the probability-overlap coefficient,

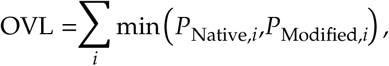

and Jensen–Shannon divergence. Projected breadth was summarized by the area within 2*RT* of the free-energy minimum and the effective number of occupied bins,

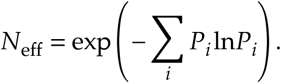

This reduced-coordinate analysis describes occupancy and exchange; it was not interpreted as a Markov-state model, evidence of equilibrium convergence, or a complete account of the linker ensemble.

Statistical inference and robustness assessment is provided in Supplementary Method S1.

## 3. Results

### 3.1. Trp698→KYN modification produces broad multimetric restriction

Across the seven prespecified global descriptors, Trp698→KYN consistently shifted the tethered linker toward greater conformational restriction (Fig. 2A,B; Tables S1 and S2). Relative to the native linker, Schlitter entropy decreased by 17.4%, covariance trace by 55.0%, and core Cα RMSF by 36.2%, whereas the effective spring constant increased approximately 3.7-fold. These concordant changes indicate reduced conformational diversity and collective mobility together with increased effective stiffness.

**Figure 2.**
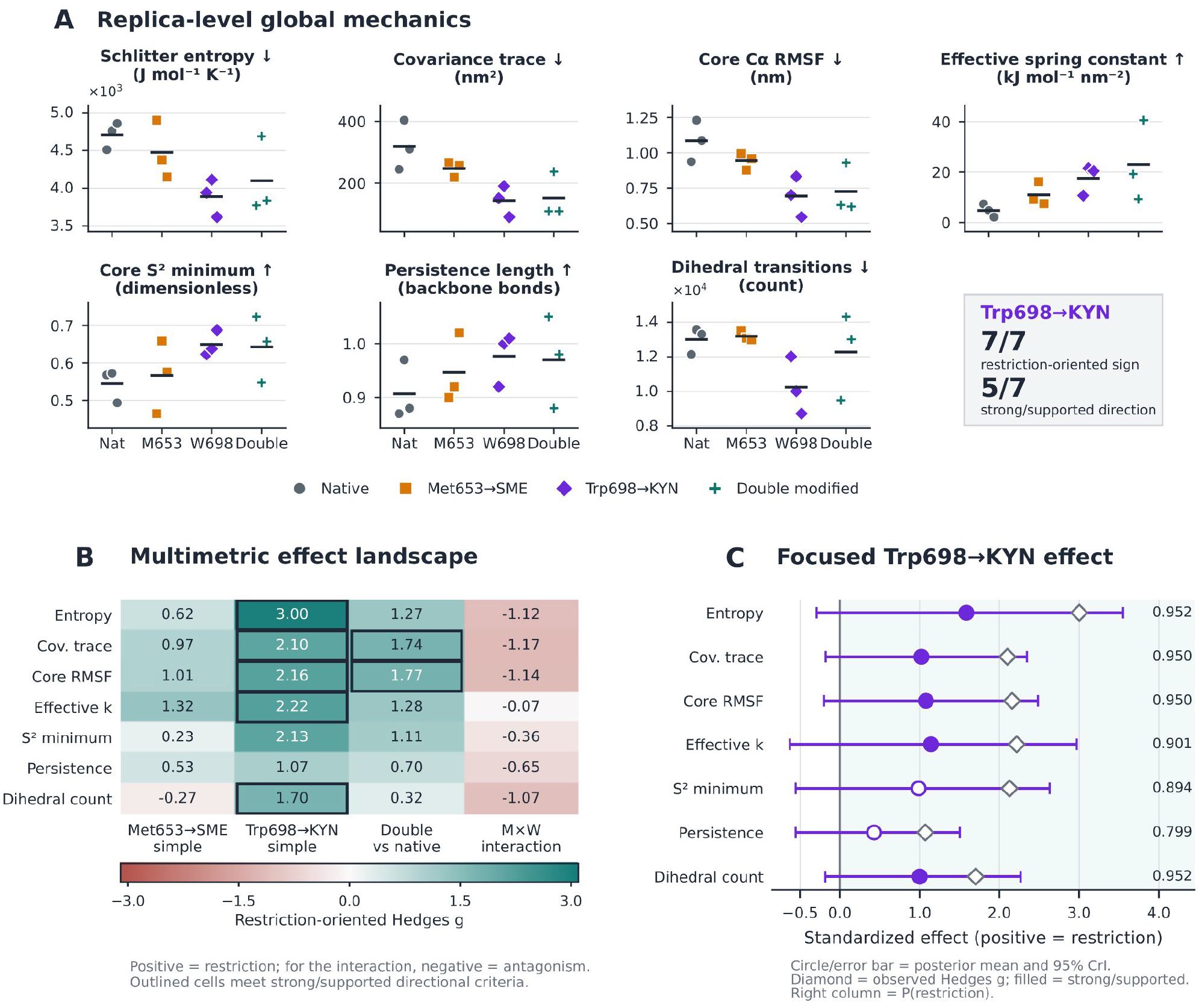
Trp698→KYN produces broad multimetric restriction of linker conformational mechanics. (A) Replica-level global descriptors for the native, Met653→SME, Trp698→KYN, and double-modified systems. Points represent independent trajectories (n=3 per condition); horizontal lines indicate replica means. Arrows denote the raw-value direction corresponding to restriction: decreased Schlitter entropy, covariance trace, core Cα RMSF, and dihedral-transition count, and increased effective spring constant, core S2min, and persistence length. (B) Restriction-oriented Hedges g for the Met653→SME and Trp698→KYN simple contrasts, double-modified versus native contrast, and Met653×Trp698 interaction. Positive values denote greater restriction; for the interaction, positive and negative values denote cooperative and antagonistic deviation from additivity, respectively. Outlined cells satisfy the prespecified strong or supported directional criteria. (C) Focused Trp698→KYN analysis. Circles and horizontal lines show posterior mean effects and 95% credible intervals on a standardized effect scale; diamonds show observed restriction-oriented Hedges g. Filled circles denote strong or supported effects; values at right give the posterior probability of restriction, Prestriction.

Local backbone order and conformational exchange changed in the same direction: core 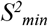 and persistence length increased, whereas the dihedral-transition count decreased by 21.2%. Restriction-oriented Hedges g was positive for all seven descriptors (1.07–3.00). Posterior analysis provided strong directional evidence for decreases in entropy, covariance trace, core RMSF, and dihedral transitions, and supported evidence for increased effective stiffness; changes in 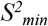 and persistence length remained exploratory. Thus, Five descriptors met the prespecified thresholds for high or supported directional probability and deletion stability (Table S2).

The 95% credible intervals for individual descriptors included zero, consistent with the limited number of independent trajectories (n=3 per condition). Inference therefore rests on multimetric convergence and directional robustness rather than any single univariate comparison. Importantly, the restriction-oriented signs of all seven Trp698→KYN effects were retained under both one-at-a-time and balanced replica-deletion analyses.

Met653→SME alone produced smaller or more heterogeneous effects, and no global descriptor met the strong or supported criteria (Fig. 2B; Table S2). The double-modified linker remained restriction-oriented relative to native across all seven descriptors, with supported reductions in covariance trace and core RMSF. Although the global factorial interactions were predominantly antagonistic in direction, none reached the supported threshold. The global analysis therefore identifies Trp698→KYN as the dominant restrictive perturbation but does not establish generalized antagonism by concurrent Met653 modification.

Supporting geometric descriptors were consistent with compaction (Fig. S1; Tables S1 and S2). Trp698→KYN decreased the radius of gyration by 23.3%, produced a smaller reduction in end-to-end distance, and increased the mean number of intramolecular hydrogen bonds from 32.5 to 45.2. These changes are consistent with a compact ensemble containing a denser internal hydrogen-bond network, although they do not establish that hydrogen-bond formation caused the restriction.

### 3.2. Restriction is spatially distributed along the linker

Residue-resolved analysis showed that Trp698→KYN-associated restriction extended well beyond the modified site (Fig. 3A). Mean Cα RMSF decreased across much of residues 630–709, with pronounced differences in the N-terminal region, residues 655–662, and the extended region spanning approximately residues 675–694. Mobility also decreased through much of the C-terminal segment containing Trp698. The double-modified condition showed a similarly broad pattern, whereas Met653→SME alone produced a smaller and more heterogeneous response.

**Figure 3.**
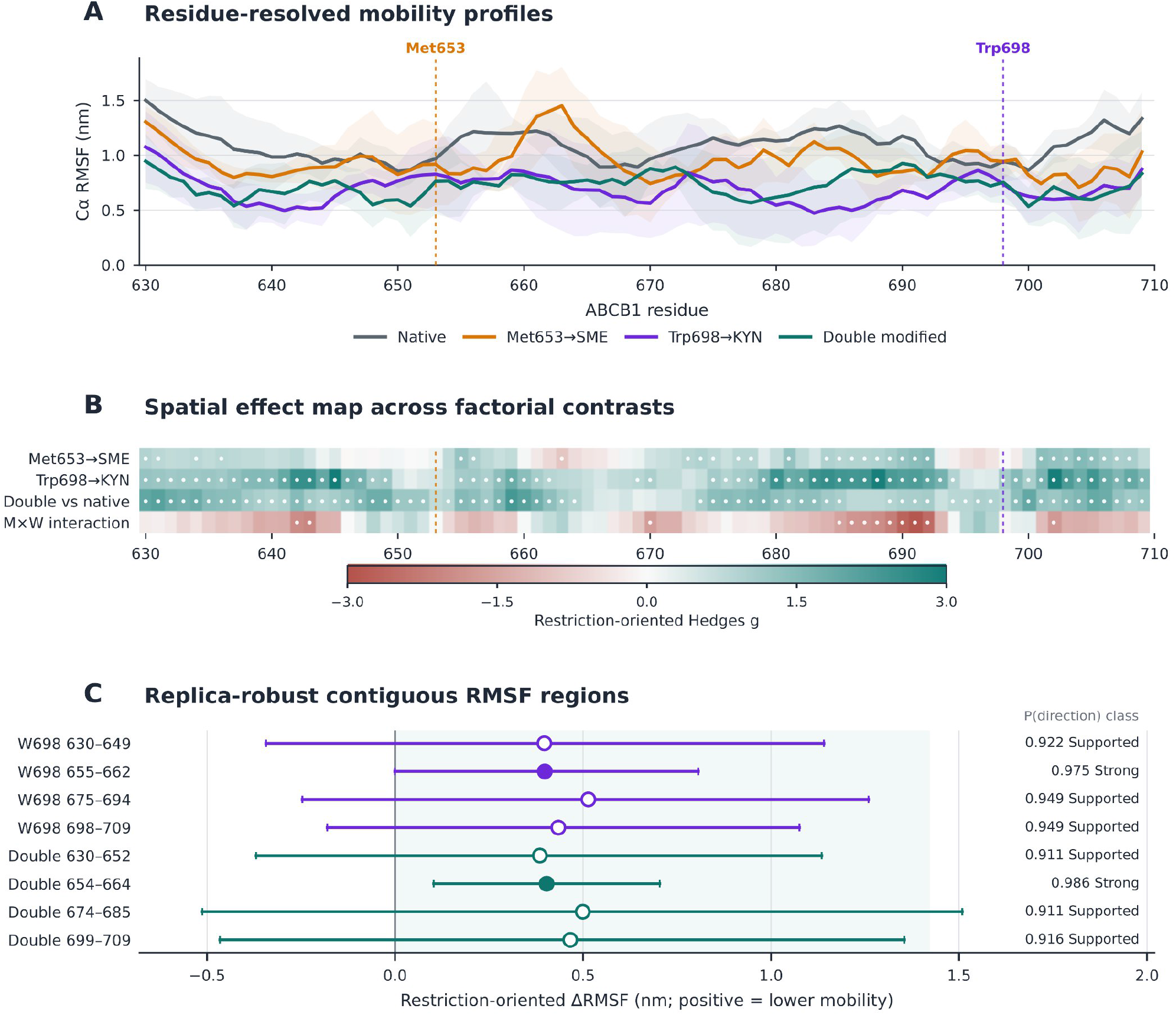
Trp698→KYN is associated with distributed and regionally coherent remodeling of linker mobility. (A) Residue-resolved Cα RMSF profiles across residues 630–709. Lines indicate condition means; faint envelopes show between-replica SDs. Vertical guides mark Met653 and Trp698. (B) Restriction-oriented RMSF effect sizes for the Met653→SME and Trp698→KYN simple contrasts, double-modified versus native contrast, and factorial interaction. Positive values denote reduced RMSF; negative interaction values denote antagonism relative to additivity. White dots identify residues meeting the candidate-residue rule. No isolated residue survived Benjamini–Hochberg false-discovery-rate correction; effects were therefore interpreted through contiguous, robustness-supported regions. (C) Replica-robust contiguous RMSF regions for the Trp698→KYN and double-modified contrasts. Points and intervals show posterior mean restriction-oriented RMSF changes and 95% credible intervals. Values at right report directional probability and robustness classification. All regions retained 100% sign stability under balanced deletion.

The spatial effect map was predominantly restriction-oriented for both the Trp698→KYN and double-modified contrasts (Fig. 3B; Table S3). Met653→SME produced fewer candidate residues, grouped into exploratory regions at 673–675, 684–692, and 701–708. The factorial interaction contained one exploratory antagonistic region at 685–692, providing no evidence for uniform antagonism along the linker.

Because no isolated residue survived false-discovery-rate correction, inference was restricted to contiguous regions satisfying the prespecified effect-size, directional-probability, and deletion-stability criteria. Four regions supported Trp698→KYN-associated restriction: residues 630–649, 655–662, 675–694, and 698–709 (Fig. 3C; Table S3). The largest standardized regional effect occurred at residues 655–662 (ΔRMSF=0.398 nm; g=2.88), more than 35 residues from the modified position, although its credible interval marginally included zero. The double-modified linker retained a comparable but non-identical pattern, with supported restriction at residues 630–652, 654–664, 674–685, and 699–709; the interval for residues 654–664 excluded zero. Every displayed Trp698→KYN and double-modification region retained its restriction-oriented sign in the deletion analyses.

Integration of RMSF with 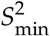 identified regions in which reduced mobility coincided with increased backbone order (Fig. S2; Table S4A). Trp698→KYN produced two consensus regions, at residues 635–643 and 676–682. The double-modified condition contained shorter consensus regions at 632–633, 646–648, and 657–658, whereas Met653→SME alone produced one short region at 673–674. The predefined Trp698-centered window (residues 696–700) did not meet the two-metric consensus criterion (Table S4B): RMSF shifted moderately toward restriction, whereas *S*^*2*^_*min*_ changed weakly in the opposite direction. Thus, the most coherent residue-level responses occurred away from the modified position, indicating distributed rather than exclusively site-centered remodeling.

### 3.3 Trp698→KYN confines the linker to a compact, low-exchange projected ensemble

Projection onto radius of gyration and end-to-end distance (*R*_*g*_ ,*R*) revealed pronounced ensemble redistribution after Trp698→KYN (Fig. 4A–D; Table S5). The native linker occupied compact, intermediate, and extended regions, with 46.9% of the ensemble assigned to the compact state (Fig. 4A,E). Met653→SME retained access to multiple regions but showed substantial trajectory-level heterogeneity (Fig. 4B). In contrast, Trp698→KYN populated a narrow, low-*R*_*g*_ basin corresponding almost exclusively to the native-defined compact state (Fig. 4C).

**Figure 4.**
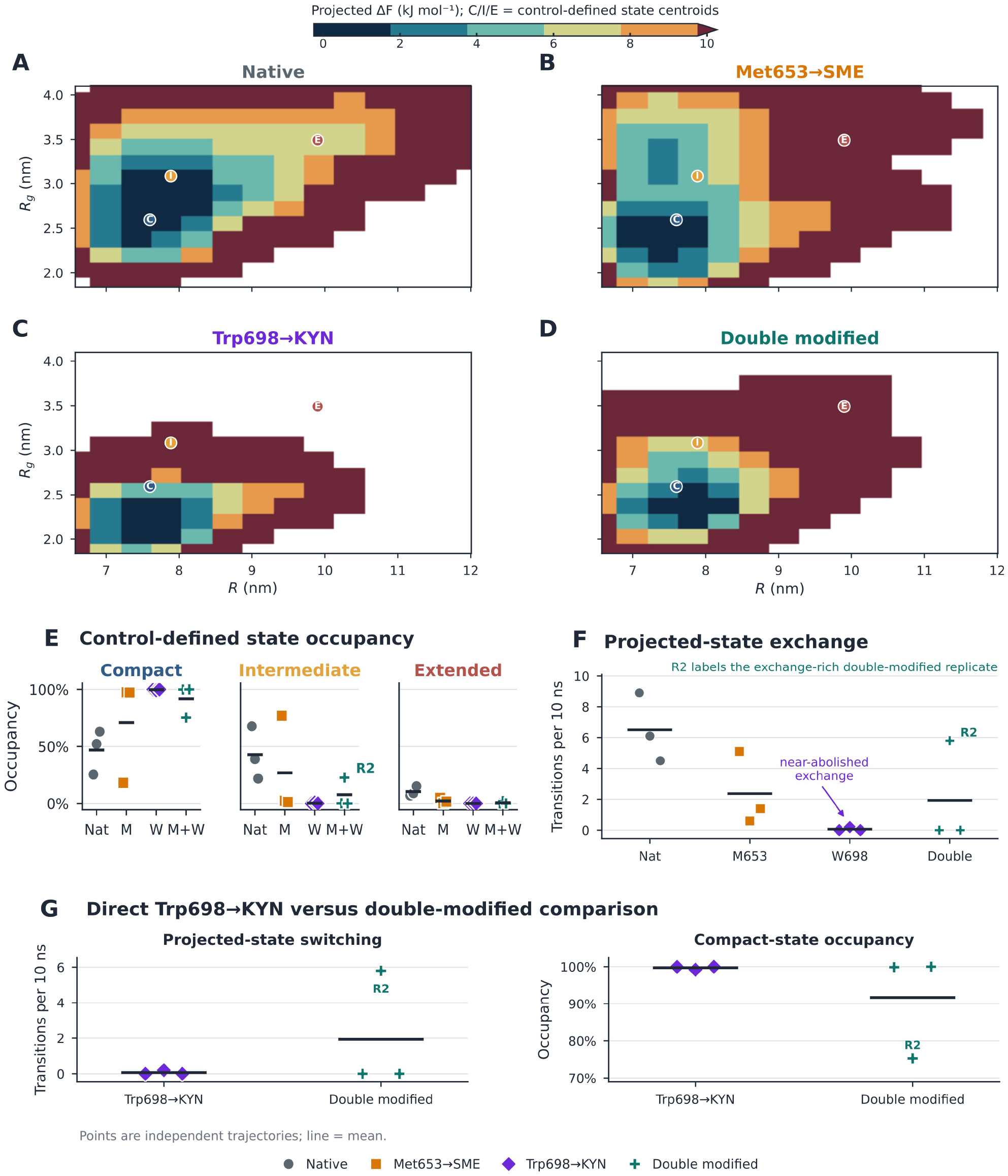
Trp698→KYN produces a compact, low-exchange projected ensemble. (A–D) Projected conformational landscapes for the (A) native, (B) Met653→SME, (C) Trp698→KYN, and (D) double-modified linker ensembles as functions of end-to-end distance (R) and radius of gyration (R_g_). All landscapes share coordinate limits and a projected ΔF scale of 0–10 kJ mol^−1^; replica histograms were normalized individually before condition averaging. C, I, and E mark native-defined Compact, Intermediate, and Extended centroids. The projections describe occupancy in the selected coordinates and are not complete free-energy surfaces or Markov-state models. (E) Replica-level occupancies of the three projected states. (F) Transitions among native-defined states per 10 ns. R2 identifies the exchange-rich double-modified replicate. (G) Direct replica-level comparison of the Trp698→KYN and double-modified conditions for projected-state switching and compact-state occupancy. Points represent independent trajectories (n=3 per condition); horizontal lines show means, and replica labels identify individual trajectories without implying pairing between conditions. The comparison is descriptive and does not invoke an arithmetic-additive baseline.

Trp698→KYN increased compact-state occupancy from 46.9% to 99.7% (bootstrap 95% CI for the change, 36.8–74.3 percentage points; Fig. 4E; Tables S5 and S7), with corresponding losses of the intermediate and extended populations. The native and modified projected distributions showed limited overlap and substantial divergence (Table S8A). Compact occupancy remained at least 98.4% in both halves of all three trajectories, supporting the temporal stability of the redistribution within the analyzed windows (Table S8B). State switching decreased from 6.50 to 0.07 transitions per 10 ns (95% CI for the change, -8.83 to -4.45; Fig. 4F; Table S7), with two modified trajectories showing no transitions. Projected ensemble breadth decreased concurrently: the effective number of occupied bins, *N*_eff_, fell by 61.1%, and the area within 2RT of the projected free-energy minimum decreased from 1.72 to 0.80 nm^2^ (Fig. S3; Tables S5 and S7). Both breadth measures retained their direction in all leave-one-replica-out analyses. Together, these findings define an almost exclusively compact, low-breadth, and low-exchange ensemble within the selected projection.

### 3.4. The double-modified condition remains restricted but shows trajectory-level heterogeneity

Met653→SME alone did not produce a uniform state redistribution. Mean compact occupancy increased to 70.9%, but the individual replica values ranged from 18.2% to 97.5% (Fig. 4B,E; Tables S5 and S6). Mean switching nevertheless decreased to 2.37 transitions per 10 ns (Fig. 4F). Met653→SME therefore reduced projected exchange without reproducibly selecting the same basin across trajectories.

The double-modified linker also remained more restricted than native: compact-state occupancy was 91.7%, switching occurred at 1.93 transitions per 10 ns, and *N*_eff_ remained substantially below the native value (Figs. 4D–F and S3; Tables S5 and S6). Concurrent Met653 modification therefore did not restore the native ensemble.

Compared with Trp698→KYN alone, the double-modified condition showed greater projected-state exchange. Mean switching increased from 0.07 to 1.93 transitions per 10 ns, while mean compact-state occupancy decreased from 99.7% to 91.7% (Fig. 4G; Tables S5 and S6). Extended-state occupancy increased only slightly, from 0.01% to 0.67% (Table S5). As shown by the replica-level distributions in Fig. 4G, this difference was strongly heterogeneous. Double-modified replicas 1 and 3 remained almost completely compact and showed no switching, whereas replica 2 switched 5.8 times per 10 ns and had a compact-state occupancy of 75.3% (Table S6). This exchange-rich trajectory also sampled a broader projected region and was retained as part of the observed ensemble rather than classified as an outlier.

Because the raw-scale additive predictions for switching and extended-state occupancy fell below their zero bounds, the corresponding factorial contrasts were not interpreted as evidence of non-additivity. Figure 4G therefore presents direct descriptive comparisons between the Trp698→KYN and double-modified conditions. The higher mean exchange in the double-modified condition was concentrated in one of three trajectories and does not establish reproducible reopening or general reversal of Trp698→KYN-associated confinement.

## 4. Discussion

The central finding is that defined oxidative lesions at two solvent-accessible sites produced markedly different ensemble responses in the mechanically tethered ABCB1 linker. Trp698→KYN generated a coherent restriction phenotype spanning global, residue-resolved, and projected-state measurements, whereas Met653→SME produced weaker and more heterogeneous remodeling. Directional stability across leave-one-replica-out analyses supports the robustness of the Trp698→KYN response, although the limited number of trajectories and broad individual uncertainty intervals preclude treating the correlated descriptors as independent statistical evidence.

The markedly stronger remodeling associated with Trp698→KYN than with Met653→SME indicates that initial solvent accessibility alone did not predict the magnitude or coherence of the conformational response. However, lesion identity and residue position varied simultaneously; the observed contrast may therefore reflect differences in lesion chemistry, local sequence environment, or their interplay. Comparisons of alternative oxidative products at the same residue, or the same chemical modification introduced at different sites, would be required to distinguish these contributions.

Residue-resolved analysis further showed that the dominant Trp698→KYN response was distributed. The most coherent RMSF and 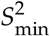 changes occurred across separated sequence regions, including segments more than 35 residues from Trp698, whereas the predefined Trp698-centered window did not meet the two-metric consensus criterion. This pattern is consistent with an intrinsically disordered region, in which a local chemical change may redistribute populations of transient contacts rather than propagate through one stable structural pathway ^[32–34]^. The results support distributed ensemble redistribution but do not identify a unique allosteric route. Likewise, the associated compaction and increased hydrogen-bond count provide a structural correlate, not evidence that hydrogen-bond formation initiates restriction.

Distributed restriction may be functionally relevant because the linker connects NBD1 to TMD2 at a position capable of influencing communication between the two halves of ABCB1. Experimental studies indicate that linker flexibility is required for coordinated ATP hydrolysis and transport ^[26]^. A lesion that reorganizes several linker regions could therefore perturb domain coupling more strongly than a narrowly local change. This remains a mechanistic prediction because neither domain coordination nor transporter activity was measured in the reduced construct.

The clearest manifestation of Trp698→KYN was the near-exclusive occupation of the native-defined compact state, accompanied by contraction of projected breadth and near elimination of state switching. This combination is more informative than a decrease in mean radius of gyration alone. Compaction specifies where the ensemble resides; reduced breadth and switching show that the ensemble also visits fewer projected configurations and exchanges less frequently among them. Trp698→KYN therefore establishes a compact, low-exchange regime rather than simply shifting the average geometry of an otherwise similarly dynamic chain. Here, “confinement” refers specifically to the (Rg,R) projection and does not imply immobilization across all microscopic degrees of freedom or equilibrium convergence. The native-defined compact, intermediate, and extended states are descriptive projection-based classes rather than a Markov-state model. Motions orthogonal to these coordinates may therefore remain active.

The projected phenotype nevertheless yields a specific functional hypothesis. ABCB1 transport depends on repeated coordination of TMD and NBD rearrangements ^[22,23,25]^, and excessive linker restriction could alter the timing or accessible range of these motions. The present data cannot determine whether the compact ensemble would inhibit ATP hydrolysis, uncouple hydrolysis from transport, alter substrate specificity, or stabilize a particular full-length state. The defensible prediction is therefore that Trp698→KYN formation may alter full-length conformational coupling, not that it necessarily suppresses drug efflux.

Met653→SME did not select a uniform projected state, and the double-modified condition remained substantially more restricted than native. Relative to Trp698→KYN alone, the double-modified condition showed higher mean switching and slightly greater access to noncompact states, but this difference arose primarily from one of three trajectories (Fig. 4G). Because the arithmetic additive predictions for the bounded switching and occupancy outcomes fell below zero, their raw-scale factorial contrasts were not treated as evidence of non-additivity. The results therefore show that an exchange-accessible projected substate was sampled in one double-modified simulation, not that Met653→SME reproducibly rescued or antagonized the Trp698→KYN phenotype. Longer simulations and additional independent replicas are needed to distinguish competing long-lived substates from incomplete sampling. One provisional explanation is that Trp698→KYN favors transient contacts that stabilize compact states, whereas Met653→SME may alter local solvation or contact preferences and thereby affect the probability of leaving this ensemble. Because contact identities, lifetimes, and residue-specific solvation were not quantified, this interpretation remains a hypothesis.

The present findings narrow the proposed CAP–ABCB1–MDR pathway to a specific, experimentally testable intermediate: defined oxidative lesions can redistribute conformational occupancy and exchange within a mechanically constrained ABCB1 linker. If such lesions form in CAP-exposed ABCB1, the resulting linker remodeling could perturb NBD–TMD coordination and thereby influence ATP hydrolysis, drug efflux, intracellular drug accumulation, and chemotherapy response. The simulations directly support only lesion-associated remodeling of the reduced linker ensemble. They do not establish that CAP generates Trp698→KYN or Met653→SME in cellular ABCB1, that these modifications alter transporter function, or that P-gp is a principal mediator of CAP-induced sensitization in resistant cells. These downstream steps therefore remain hypotheses requiring validation in full-length ABCB1 and cellular models. This evidentiary boundary makes the hypothesis experimentally tractable. Targeted LC–MS/MS of ABCB1 isolated before and after CAP or plasma-activated-medium exposure should determine whether kynurenine formation at Trp698, methionine sulfoxide formation at Met653, or other linker lesions form and at what occupancy. Time-resolved measurements could establish whether modification precedes functional change. ATPase activity, fluorescent-substrate or chemotherapeutic efflux, and intracellular drug accumulation should then be quantified while independently controlling for ABCB1 abundance, membrane localization, viability, and nonspecific membrane damage. Chemically defined incorporation of kynurenine or stereochemically characterized methionine sulfoxide would offer a more specific test than conventional amino-acid substitutions.

Full-length, membrane-embedded simulations provide a complementary next step. Introducing the modifications into multiple inward-facing and nucleotide-bound ABCB1 structures would allow direct evaluation of NBD separation and dimerization, coupling-helix contacts, linker–domain interactions, substrate-access pathways, and transport-cycle transitions. Multiple starting structures, longer sampling, and additional replicas will be required to determine whether the compact linker phenotype persists in the presence of the domains and membrane. Several limitations constrain interpretation. The tethered construct approximates the terminal separation imposed by neighboring domains but omits the TMDs, NBDs, intracellular loops, membrane, nucleotide, and substrate. KYN and SME were represented as fixed classical-force-field residues. The simulations do not describe lesion formation, reaction rates, occupancies, repair, or competition with other oxidative products. Only two sites and one product per site were examined, whereas CAP-associated chemistry may generate multiple side-chain and backbone modifications. SASA identifies contrasting exposure regimes but not oxidation susceptibility in full-length ABCB1. Projected occupancies, breadth, and switching depend on the selected coordinates, grid, native-derived clustering, and filtering procedure and are not microscopic kinetic constants or proof of equilibrium. These limitations make direct modification measurements, full-length simulations, and functional validation essential.

## 5. Conclusions

Trp698→KYN produced a broad, directionally concordant restriction phenotype in the mechanically tethered ABCB1 linker. Remodeling extended across separated sequence regions and culminated in an almost exclusively compact, low-breadth, and low-exchange projected ensemble. Met653→SME produced a weaker and more heterogeneous response. Although the double-modified condition showed greater mean projected exchange than Trp698→KYN alone, this behavior was concentrated in one of three trajectories and therefore does not establish reproducible restoration, antagonism, or non-additive counteraction. These findings identify lesion-associated remodeling of the ABCB1 linker as a plausible intermediate between oxidative chemistry and altered transporter regulation. They do not establish lesion formation after CAP exposure or consequent changes in drug transport. The critical next steps are therefore to detect and quantify these lesions in CAP-exposed ABCB1 and determine whether they alter conformational coupling, ATPase activity, drug efflux, and chemotherapy response in the full-length transporter.

## Declarations

**CRediT** authorship contribution statement. Mansoureh Hoseini: Conceptualization, Methodology, Software, Formal analysis, Investigation, Data curation, Visualization, Writing – original draft. Hossein Abbasi: Supervision, Conceptualization, Methodology, Writing – review & editing.

## Data availability

All tabulated replica-level results are provided in Source_Data.csv and Supplementary Tables S1–S8. All 12 trajectories (200 ns each; 2.4 μs total; no plasma species; SME/KYN lesions) were generated de novo for this study and no trajectories were reused from other work. Input topologies and analysis scripts are available from the corresponding author on reasonable request.

## References

1. Bukowski, K., Kciuk, M., & Kontek, R. (2020). Mechanisms of multidrug resistance in cancer chemotherapy. International Journal of Molecular Sciences, 21(9), 3233. 10.3390/ijms21093233

2. Karthika, C., Sureshkumar, R., Zehravi, M., Akter, R., Ali, F., Ramproshad, S., Mondal, B., Tagde, P., Ahmed, Z., Khan, F. S., & others. (2022). Multidrug resistance of cancer cells and the vital role of P-glycoprotein. Life, 12(6), 897. 10.3390/life12060897

3. Seelig, A. (2020). P-glycoprotein: One mechanism, many tasks and the consequences for pharmacotherapy of cancers. Frontiers in Oncology, 10, 576559. 10.3389/fonc.2020.576559

4. Gottesman, M. M., Lavi, O., Hall, M. D., & Gillet, J.-P. (2016). Toward a better understanding of the complexity of cancer drug resistance. Annual Review of Pharmacology and Toxicology, 56(1), 85–102. 10.1146/annurev-pharmtox-010715-103111

5. Tia, S. T., Luo, M., & Fan, W. (2025). Mapping the role of P-gp in multidrug resistance: Insights from recent structural studies. International Journal of Molecular Sciences, 26(9), 4179. 10.3390/ijms26094179

6. Al Thubiani, W. S. (2023). The role of P-glycoprotein (P-gp) in cancer multidrug resistance (MDR): Challenges for inhibiting P-gp in the context of overcoming MDR. Journal of Pharmaceutical Research International, 35(23), 44–58. 10.9734/jpri/2023/v35i237422

7. Szakács, G., Paterson, J. K., Ludwig, J. A., Booth-Genthe, C., & Gottesman, M. M. (2006). Targeting multidrug resistance in cancer. Nature Reviews Drug Discovery, 5(3), 219–234. 10.1038/nrd1984

8. Hoosain, F. G., Choonara, Y. E., Tomar, L. K., Kumar, P., Tyagi, C., du Toit, L. C., & Pillay, V. (2015). Bypassing P-glycoprotein drug efflux mechanisms: Possible applications in Pharmacoresistant schizophrenia therapy. BioMed Research International, 2015(1), 484963. 10.1155/2015/484963

9. Dei, S., Braconi, L., Romanelli, M. N., & Teodori, E. (2019). Recent advances in the search of BCRP-and dual P-gp/BCRP-based multidrug resistance modulators. Cancer Drug Resistance, 2(3), 710. 10.20517/cdr.2019.31

10. Robey, R. W., Pluchino, K. M., Hall, M. D., Fojo, A. T., Bates, S. E., & Gottesman, M. M. (2018). Revisiting the role of ABC transporters in multidrug-resistant cancer. Nature Reviews Cancer, 18(7), 452–464. 10.1038/s41568-018-0005-8

11. Yan, D., Talbot, A., Nourmohammadi, N., Cheng, X., Canady, J., Sherman, J., & Keidar, M. (2015). Principles of using cold atmospheric plasma stimulated media for cancer treatment. Scientific Reports, 5(1), 18339. 10.1038/srep18339

12. Yan, D., Sherman, J. H., & Keidar, M. (2016). Cold atmospheric plasma, a novel promising anti-cancer treatment modality. Oncotarget, 8(9), 15977. 10.18632/oncotarget.13304

13. Keidar, M., Walk, R., Shashurin, A., Srinivasan, P., Sandler, A., Dasgupta, S., Ravi, R., Guerrero-Preston, R., & Trink, B. (2011). Cold plasma selectivity and the possibility of a paradigm shift in cancer therapy. British Journal of Cancer, 105(9), 1295–1301. 10.1038/bjc.2011.386.

14. Keidar, M., Yan, D., & Lin, L. (2023). Cold atmospheric plasma-based cancer therapy. IOP Publishing.

15. Murillo, D., Huergo, C., Gallego, B., Rodríguez, R., & Tornín, J. (2023). Exploring the use of cold atmospheric plasma to overcome drug resistance in cancer. Biomedicines, 11(1), 208. 10.3390/biomedicines11010208

16. Pefani-Antimisiari, K., Athanasopoulos, D. K., Marazioti, A., Sklias, K., Rodi, M., De Lastic, A.-L., Mouzaki, A., Svarnas, P., & Antimisiaris, S. G. (2021). Synergistic effect of cold atmospheric pressure plasma and free or liposomal doxorubicin on melanoma cells. Scientific Reports, 11(1), 14788. 10.1038/s41598-021-94130-7

17. Park, S., Kim, H., Ji, H. W., Kim, H. W., Yun, S. H., Choi, E. H., & Kim, S. J. (2019). Cold atmospheric plasma restores paclitaxel sensitivity to paclitaxel-resistant breast cancer cells by reversing expression of resistance-related genes. Cancers, 11(12), 2011. 10.3390/cancers11122011

18. Nitsch, A., Qarqash, S., Römer, S., Schoon, J., Singer, D., Bekeschus, S., Ekkernkamp, A., Wassilew, G. I., Tzvetkov, M. V., & Haralambiev, L. (2024). Effective combination of cold physical plasma and chemotherapy against Ewing sarcoma cells in vitro. Scientific Reports, 14(1), 6505. 10.1038/s41598-024-56985-4

19. Mihai, C.-T., Mihaila, I., Pasare, M. A., Pintilie, R. M., Ciorpac, M., & Topala, I. (2022). Cold atmospheric plasma-activated media improve paclitaxel efficacy on breast cancer cells in a combined treatment model. Current Issues in Molecular Biology, 44(5), 1995–2014. 10.3390/cimb44050135

20. Aller SG et al. (2009). Structure of P-glycoprotein reveals a molecular basis for poly-specific drug binding. Science, 323(5922):1718–1722. 10.1126/science.1168750

21. Kim, Y., & Chen, J. (2018). Molecular structure of human P-glycoprotein in the ATP-bound, outward-facing conformation. Science, 359(6378), 915–919. 10.1126/science.aar7389

22. Ward, A. B., Szewczyk, P., Grimard, V., Lee, C.-W., Martinez, L., Doshi, R., Caya, A., Villaluz, M., Pardon, E., Cregger, C., & others. (2013). Structures of P-glycoprotein reveal its conformational flexibility and an epitope on the nucleotide-binding domain. Proceedings of the National Academy of Sciences, 110(33), 13386–13391. 10.1073/pnas.1309275110

23. Rani, P., Mandal, P., Rajak, B. K., & Singh, D. V. (2024). A review on dynamics of permeability-glycoprotein in efflux of chemotherapeutic drugs. Frontiers in Drug Discovery, 4, 1363364. 10.3389/fddsv.2024.1363364

24. Sharom, F. J. (2014). Complex interplay between the P-glycoprotein multidrug efflux pump and the membrane: Its role in modulating protein function. Frontiers in Oncology, 4, 41. 10.3389/fonc.2014.00041

25. Li, Y., Yuan, H., Yang, K., Xu, W., Tang, W., & Li, X. (2010). The structure and functions of P-glycoprotein. Current Medicinal Chemistry, 17(8), 786–800. 10.2174/092986710790514507

26. Hrycyna, C. A., Airan, L. E., Germann, U. A., Ambudkar, S. V., Pastan, I., & Gottesman, M. M. (1998). Structural flexibility of the linker region of human P-glycoprotein permits ATP hydrolysis and drug transport. Biochemistry, 37(39), 13660–13673. 10.1021/bi9808823

27. Sato, T., Kodan, A., Kimura, Y., Ueda, K., Nakatsu, T., & Kato, H. (2009). Functional role of the linker region in purified human P-glycoprotein. The FEBS Journal, 276(13), 3504–3516. 10.1111/j.1742-4658.2009.07072.x

28. Ferreira, R. J., Ferreira, M.-J. U., & Dos Santos, D. J. (2012). Insights on P-glycoprotein’s efflux mechanism obtained by molecular dynamics simulations. Journal of Chemical Theory and Computation, 8(6), 1853–1864. 10.1021/ct300083m

29. Ferreira, R. J., Ferreira, M.-J. U., & Dos Santos, D. J. (2013). Assessing the stabilization of P-glycoprotein’s nucleotide-binding domains by the linker, using molecular dynamics. Molecular Informatics, 32(5–6), 529–540. 10.1002/minf.201200175

30. Elbahnsi, A., Dudas, B., Cisternino, S., Declèves, X., & Miteva, M. A. (2024). Mechanistic insights into P-glycoprotein ligand transport and inhibition revealed by enhanced molecular dynamics simulations. Computational and Structural Biotechnology Journal, 23, 2548–2564. 10.1016/j.csbj.2024.06.010

31. Han, S. B., Warwicker, J., Fan, H., & Prince, S. M. (2025). Nucleotide asymmetry and flexible linker dynamics modulate drug efflux cycle of P-glycoprotein, A computational study. Computational and Structural Biotechnology Journal, 27, 4825–4837. 10.1016/j.csbj.2025.10.064

32. Thomasen, F. E., & Lindorff-Larsen, K. (2022). Conformational ensembles of intrinsically disordered proteins and flexible multidomain proteins. Biochemical Society Transactions, 50(1), 541–554. 10.1042/BST20210499

33. Sørensen, C. S., & Kjaergaard, M. (2019). Effective concentrations enforced by intrinsically disordered linkers are governed by polymer physics. Proceedings of the National Academy of Sciences, 116(46), 23124–23131. 10.1073/pnas.1904813116

34. Sulc, J., & Vondrasek, J. (2025). Unraveling the tether: Exploring representative protein linkers and their structural and thermodynamical properties. The Journal of Physical Chemistry B, 129(15), 3720–3730. 10.1021/acs.jpcb.4c04194

35. Hoseini, M., & Abbasi, H. (2025). Selected plasma-associated molecular species remodel the ABCB1 linker with effects retained under structure-derived tethering. bioRxiv 2025.08.13.670033. 10.1101/2025.08.13.670033

36. Hoseini, M., & Abbasi, H. (2026). Composition-Dependent Remodeling of the ABCB1 Disordered Linker by Plasma-Generated Reactive Mixtures. ChemRxiv 15010176. 10.26434/chemrxiv.15010176

37. Lewis, A. K., Dunleavy, K. M., Senkow, T. L., Her, C., Horn, B. T., Jersett, M. A., Mahling, R., McCarthy, M. R., Perell, G. T., Valley, C. C., & others. (2016). Oxidation increases the strength of the methionine-aromatic interaction. Nature Chemical Biology, 12(10), 860–866. 10.1038/nchembio.2159

38. Ehrenshaft, M., Deterding, L. J., & Mason, R. P. (2015). Tripping up Trp: Modification of protein tryptophan residues by reactive oxygen species, modes of detection, and biological consequences. Free Radical Biology and Medicine, 89, 220–228. 10.1016/j.freeradbiomed.2015.08.003

39. Jumper, J., Evans, R., Pritzel, A., Green, T., Figurnov, M., Ronneberger, O., Tunyasuvunakool, K., Bates, R., Žídek, A., Potapenko, A., & others. (2021). Highly accurate protein structure prediction with AlphaFold. Nature, 596(7873), 583–589. 10.1038/s41586-021-03819-2

40. Jo, S., Kim, T., Iyer, V. G., & Im, W. (2008). CHARMM-GUI: a web-based graphical user interface for CHARMM. Journal of Computational Chemistry, 29(11), 1859–1865.

41. Abraham, M. J., Murtola, T., Schulz, R., Páll, S., Smith, J. C., Hess, B., & Lindahl, E. (2015). GROMACS: High performance molecular simulations through multi-level parallelism from laptops to supercomputers. SoftwareX, 1, 19–25. 10.1016/j.softx.2015.06.001

42. Huang, J., Rauscher, S., Nawrocki, G., Ran, T., Feig, M., De Groot, B. L., Grubmüller, H., & MacKerell Jr, A. D. (2017). CHARMM36m: An improved force field for folded and intrinsically disordered proteins. Nature Methods, 14(1), 71–73. 10.1038/nmeth.4067

