## Supplementary Information for "Modeled Oxidative Lesions Remodel the ABCB1 Linker: A Hypothesis for Transporter Regulation"

*for*

* Corresponding author.

Corresponding Author's Information:

Hossein Abbasi

### Supplementary methods

#### Supplementary Method S1: Statistical inference and robustness assessment

All condition-level inference used replica summaries as independent observations ($n = 3$ per condition). Frames were used to estimate within-trajectory quantities and for temporal sensitivity analyses but were not treated as independent samples. Descriptive results are reported as replica mean ± sample SD.

Global descriptors were analyzed separately using a robust Bayesian hierarchical cell-means model with a Student-$t$ likelihood (four degrees of freedom) and partial pooling across the four modification states. Models were fitted in Python using four chains, with 2,600 posterior draws retained per chain after warm-up. All reported parameters satisfied $\hat{R} \leq 1.01$. Posterior contrasts were summarized by their mean, 95% credible interval, and probability of the observed biological direction.

Small-sample-corrected Hedges $g$ was calculated from replica-level observations. Effects were oriented so that positive values consistently denoted greater restriction: signs were reversed for entropy, covariance trace, RMSF, and dihedral-transition count, whereas the original sign was retained for effective stiffness, $S^2_{min}$, and persistence length. For factorial interactions, positive oriented values represented cooperative restriction relative to additivity and negative values represented antagonistic non-additivity. These terms describe metric-level behavior within the simulation factorial and do not imply pharmacological synergy or antagonism.

Directional evidence was classified as strong when the posterior probability of the observed direction was ≥ 0.95 and at least 90% of balanced deletions retained that direction; supported at probability ≥ 0.90 with at least 80% sign retention; exploratory at probability ≥ 0.75; and uncertain below 0.75. The seven primary descriptors were treated as complementary, correlated measures rather than independent statistical tests.

Landscape contrasts used replica-balanced nonparametric resampling. Two-condition comparisons were evaluated through all 729 ordered bootstrap combinations obtainable from two sets of three replica means. Four-condition factorial interactions used 200,000 fixed-seed bootstrap draws. Robustness was further assessed by single-replica deletion and balanced deletion of one replica from each contributing condition. Occupancies, overlap, divergence, breadth, modal coordinates, and switching were recalculated over the early and late halves of the analysis interval (100–150 and 150–200 ns). A result was considered replica robust only when its qualitative direction was retained under the prespecified deletion and, where applicable, temporal-window analyses.

For bounded projected outcomes, raw-scale factorial interactions were not interpreted when the corresponding arithmetic additive expectation fell outside the physically permissible range.

#### Supplementary figures

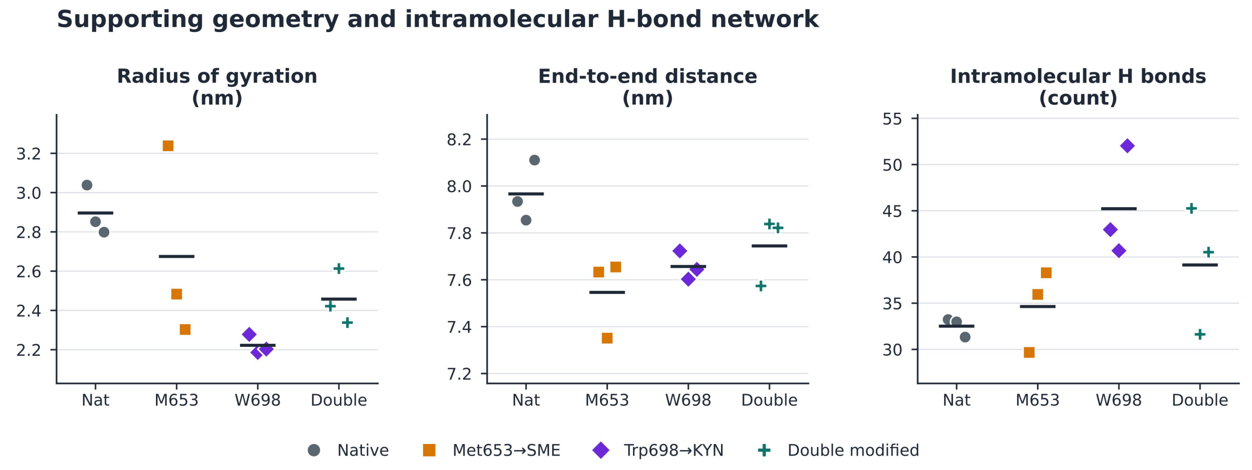


***Figure S1. Supporting changes in global linker geometry and intramolecular hydrogen bonding.*** *Replica-level radius of gyration ($R_g$), end-to-end distance, and intramolecular hydrogen-bond count calculated for the native, Met653⟶SME, Trp698⟶KYN, and double-modified linker systems. Each point represents an independent trajectory replica (n = 3 per condition), and horizontal lines indicate replica means. The geometric descriptors provide complementary evidence of modification-dependent changes in linker compaction and overall chain dimensions, while hydrogen-bond counts report accompanying changes in the intramolecular contact network.*

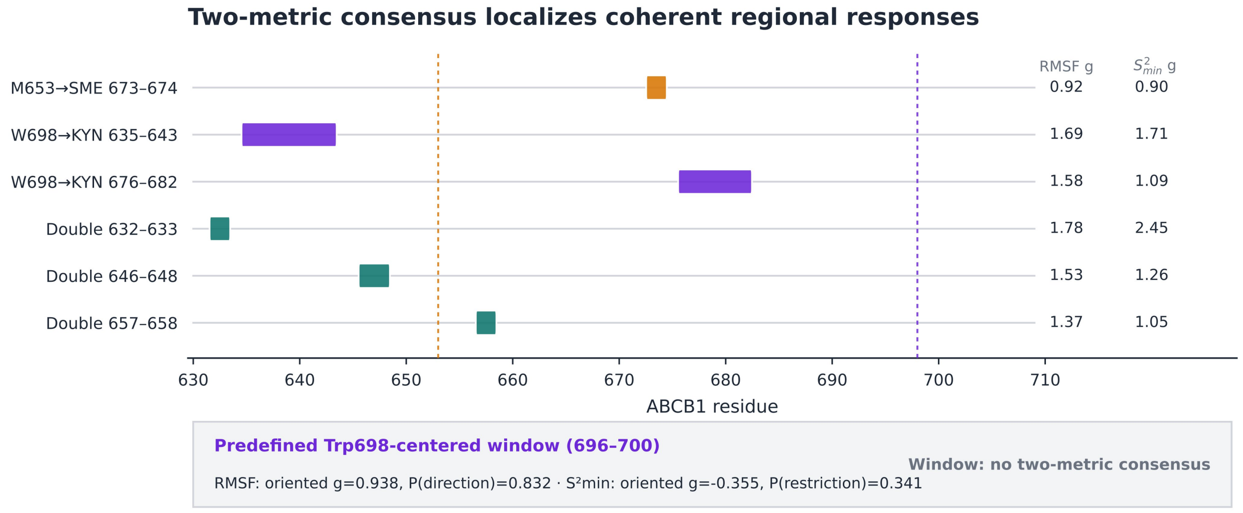


***Figure S2. Cross-metric consensus identifies spatially coherent regions of linker restriction.*** *Contiguous consensus regions across the core ABCB1 linker (residues 630–709) were defined by adjacent residues meeting the prespecified candidate criteria for both Cα RMSF and minimum $S^2$ with concordant restriction-oriented direction. Colored bands indicate consensus regions associated with the Met653⟶SME, Trp698⟶KYN, or double-modified contrasts. Values at right report the mean restriction-oriented Hedges g within each region for RMSF and $S^2_{min}$, respectively; positive values correspond to decreased positional mobility for RMSF and increased local ordering for $S^2_{min}$. Vertical guides indicate Met653 and Trp698. The predefined Trp698-centered window (residues 696–700) was additionally evaluated independently for RMSF and $S^2_{min}$. This window did not satisfy the two-metric consensus criterion, indicating that the most coherent modification-associated responses were not confined to the immediate vicinity of Trp698.*

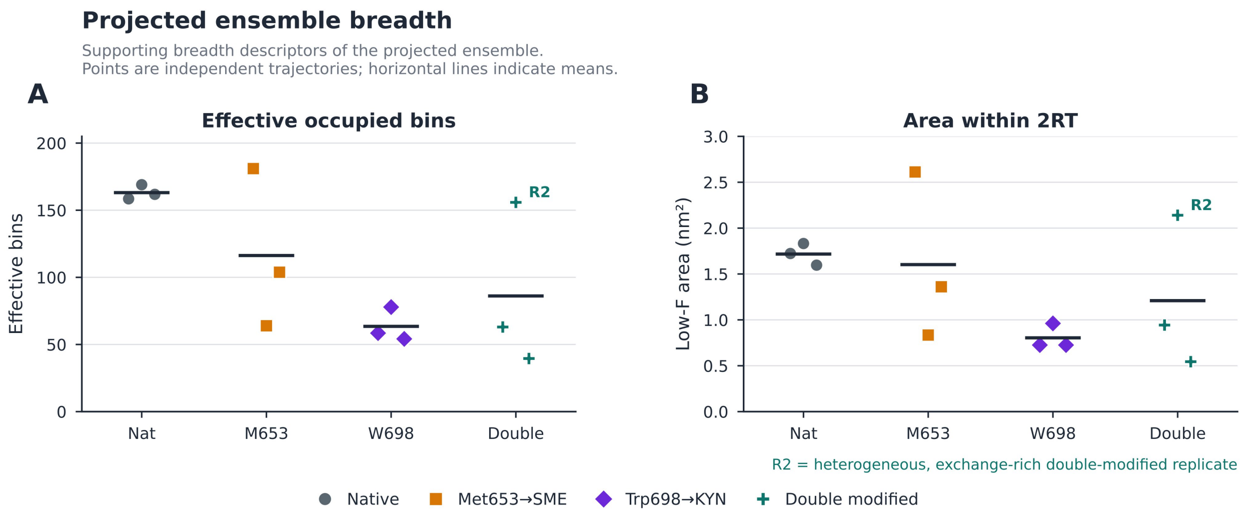


***Figure S3. Projected ensemble breadth corroborates the compact Trp698→KYN phenotype.*** *Replica-level measures of projected conformational breadth for the native, Met653→SME, Trp698→KYN, and double-modified linker ensembles. (A) Effective number of occupied bins in the radius-of-gyration/end-to-end-distance projection, providing a probability-weighted measure of how broadly each trajectory samples the projected coordinate space. (B) Area of the projected landscape lying within 2RT of the low-ΔF region, providing a complementary measure of the spatial extent of the readily populated projected ensemble. Each point represents an independent trajectory (n = 3 per condition), and horizontal lines indicate condition means. R2 identifies the heterogeneous, exchange-rich replicate of the double-modified condition.*

#### Supplementary tables

***Table S1.*** *Condition-level means for global mechanical and supporting descriptors.*

| Descriptor | Native | Met653→SME | Trp698→KYN | Double | Unit | Role |
|---|---|---|---|---|---|---|
| Schlitter entropy | 4708.6 | 4475.0 | 3890.8 | 4099.1 | $J\ mol^{-1}\ K^{-1}$ | Primary |
| Covariance trace | 319.6 | 247.6 | 143.8 | 151.7 | $nm^2$ | Primary |
| Core Cα RMSF | 1.085 | 0.943 | 0.692 | 0.726 | nm | Primary |
| Effective spring constant | 4.74 | 10.92 | 17.54 | 23.04 | $kJ\ mol^{-1}\ nm^{-2}$ | Primary |
| Core $S^2$min | 0.545 | 0.566 | 0.650 | 0.643 | dimensionless | Primary |
| Persistence length | 0.907 | 0.947 | 0.977 | 0.970 | backbone bonds | Primary |
| Dihedral-transition count | 13000 | 13189 | 10250 | 12271 | count | Primary |
| Radius of gyration (R_g) | 2.896 | 2.675 | 2.222 | 2.458 | nm | Supporting |
| End-to-end distance | 7.966 | 7.546 | 7.657 | 7.744 | nm | Supporting |
| Intramolecular H-bond count | 32.5 | 34.6 | 45.2 | 39.1 | count | Supporting |

**Note.** Values are means across three independent trajectories per condition over the 100–200 ns analysis window. Primary descriptors were prespecified for factorial inference; supporting descriptors provide geometric or contact-network context.

***Table S2.*** *Complete factorial contrast statistics for the seven prespecified global descriptors.*

| Descriptor | Contrast | Unit | Oriented observed effect | Oriented posterior mean (95% CrI) | Oriented g | P(direction) | Evidence | Deletion stability |
|---|---|---|---|---|---|---|---|---|
| Schlitter entropy | Met653→SME | J mol⁻¹ K⁻¹ | +233.7 | +33.1 (-424.8 to +514.0) | +0.62 | 0.547 | Uncertain | 89%; single stable |
| Schlitter entropy | Trp698→KYN | J mol⁻¹ K⁻¹ | +817.8 | +432.0 (-79.6 to +966.9) | +3.00 | 0.952 | Strong | 100%; single stable |
| Schlitter entropy | Double vs native | J mol⁻¹ K⁻¹ | +609.5 | +304.4 (-214.8 to +872.5) | +1.27 | 0.865 | Exploratory | 100%; single stable |
| Schlitter entropy | Met653 × Trp698 interaction | J mol⁻¹ K⁻¹ | -441.9 | -160.7 (-923.4 to +575.5) | -1.12 | 0.337 | Uncertain | 91%; single stable |
| Covariance trace | Met653→SME | nm² | +72.0 | +7.9 (-78.9 to +103.1) | +0.97 | 0.554 | Uncertain | 100%; single stable |
| Covariance trace | Trp698→KYN | nm² | +175.8 | +85.5 (-14.9 to +196.2) | +2.10 | 0.950 | Strong | 100%; single stable |
| Covariance trace | Double vs native | nm² | +167.9 | +80.6 (-22.0 to +195.6) | +1.74 | 0.931 | Supported | 100%; single stable |
| Covariance trace | Met653 × Trp698 interaction | nm² | -79.9 | -12.8 (-152.9 to +123.7) | -1.17 | 0.438 | Uncertain | 95%; single stable |
| Core Cα RMSF | Met653→SME | nm | +0.142 | +0.016 (-0.186 to +0.233) | +1.01 | 0.554 | Uncertain | 100%; single stable |
| Core Cα RMSF | Trp698→KYN | nm | +0.392 | +0.196 (-0.036 to +0.451) | +2.16 | 0.950 | Strong | 100%; single stable |
| Core Cα RMSF | Double vs native | nm | +0.359 | +0.183 (-0.053 to +0.446) | +1.77 | 0.935 | Supported | 100%; single stable |
| Core Cα RMSF | Met653 × Trp698 interaction | nm | -0.175 | -0.030 (-0.360 to +0.288) | -1.14 | 0.437 | Uncertain | 94%; single stable |
| Effective spring constant | Met653→SME | kJ mol⁻¹ nm⁻² | +6.18 | +1.62 (-7.99 to +11.19) | +1.32 | 0.636 | Uncertain | 100%; single stable |
| Effective spring constant | Trp698→KYN | kJ mol⁻¹ nm⁻² | +12.80 | +6.60 (-3.62 to +17.14) | +2.22 | 0.901 | Supported | 100%; single stable |
| Effective spring constant | Double vs native | kJ mol⁻¹ nm⁻² | +18.29 | +6.84 (-4.14 to +19.16) | +1.28 | 0.889 | Exploratory | 100%; single stable |
| Effective spring constant | Met653 × Trp698 interaction | kJ mol⁻¹ nm⁻² | -0.68 | -1.38 (-16.81 to +15.08) | -0.07 | 0.428 | Uncertain | 44% |
| Core S²min | Met653→SME | dimensionless | +0.022 | -0.003 (-0.086 to +0.077) | +0.23 | 0.471 | Uncertain | 56% |
| Core S²min | Trp698→KYN | dimensionless | +0.105 | +0.049 (-0.027 | +2.13 | 0.894 | Exploratory | 100%; |

| Descriptor | Contrast | Unit | Oriented observed effect | Oriented posterior mean (95% CrI) | Oriented g | P(direction) | Evidence | Deletion stability |
|---|---|---|---|---|---|---|---|---|
| | | | | to +0.130) | | | | single stable |
| Core S²min | Double vs native | dimensionless | +0.098 | +0.041 (-0.043 to +0.132) | +1.11 | 0.828 | Exploratory | 100%; single stable |
| Core S²min | Met653 × Trp698 interaction | dimensionless | -0.029 | -0.005 (-0.133 to +0.122) | -0.36 | 0.469 | Uncertain | 70% |
| Persistence length | Met653→SME | backbone bonds | +0.040 | +0.004 (-0.059 to +0.073) | +0.53 | 0.539 | Uncertain | 78%; single stable |
| Persistence length | Trp698→KYN | backbone bonds | +0.070 | +0.028 (-0.036 to +0.098) | +1.07 | 0.799 | Exploratory | 100%; single stable |
| Persistence length | Double vs native | backbone bonds | +0.063 | +0.021 (-0.047 to +0.094) | +0.70 | 0.718 | Uncertain | 100%; single stable |
| Persistence length | Met653 × Trp698 interaction | backbone bonds | -0.047 | -0.011 (-0.121 to +0.095) | -0.65 | 0.422 | Uncertain | 75%; single stable |
| Dihedral-transition count | Met653→SME | count | -189 | -381 (-1988 to +1213) | -0.27 | 0.305 | Uncertain | 67% |
| Dihedral-transition count | Trp698→KYN | count | +2750 | +1617 (-293 to +3662) | +1.70 | 0.952 | Strong | 100%; single stable |
| Dihedral-transition count | Double vs native | count | +729 | -28 (-1818 to +1930) | +0.32 | 0.481 | Uncertain | 67% |
| Dihedral-transition count | Met653 × Trp698 interaction | count | -1832 | -1263 (-4185 to +1652) | -1.07 | 0.195 | Exploratory | 94%; single stable |

**Note.** Effects are oriented so positive values indicate restriction for simple and total contrasts and cooperative restriction for the factorial interaction. P denotes P(restriction) for simple and total contrasts and P(cooperative restriction) for the interaction. CrI, Bayesian credible interval. Balanced-deletion stability is followed by whether all single-replica deletions retained the sign.

***Table S3.*** *Contiguous residue-level Cα RMSF regions identified by the prespecified hotspot rule.*

| Contrast | Region | Direction | Oriented ΔRMSF (nm) | Oriented posterior mean (95% CrI) | Oriented g | P(direction) | Balanced stability | Evidence |
|---|---|---|---|---|---|---|---|---|
| Met653→SME | 673–675 | Restriction | +0.226 | +0.224 (-0.481 to +0.919) | +0.96 | 0.833 | 100% | Exploratory |
| Met653→SME | 684–692 | Restriction | +0.244 | +0.242 (-0.372 to +0.830) | +1.24 | 0.878 | 100% | Exploratory |
| Met653→SME | 701–708 | Restriction | +0.353 | +0.353 (-0.471 to +1.169) | +1.30 | 0.891 | 100% | Exploratory |
| Trp698→KYN | 630–649 | Restriction | +0.399 | +0.397 (-0.343 to +1.141) | +1.63 | 0.922 | 100% | Supported |
| Trp698→KYN | 655–662 | Restriction | +0.396 | +0.398 (-0.001 to +0.807) | +2.88 | 0.975 | 100% | Strong |
| Trp698→KYN | 675–694 | Restriction | +0.519 | +0.514 (-0.247 to +1.259) | +2.07 | 0.949 | 100% | Supported |
| Trp698→KYN | 698–709 | Restriction | +0.435 | +0.434 (-0.180 to +1.075) | +2.08 | 0.949 | 100% | Supported |
| Double vs native | 630–652 | Restriction | +0.383 | +0.385 (-0.369 to +1.135) | +1.53 | 0.911 | 100% | Supported |
| Double vs native | 654–664 | Restriction | +0.403 | +0.403 (+0.103 to +0.704) | +4.07 | 0.986 | 100% | Strong |
| Double vs native | 674–685 | Restriction | +0.500 | +0.499 (-0.513 to +1.509) | +1.40 | 0.911 | 100% | Supported |
| Double vs native | 689–692 | Restriction | +0.229 | +0.227 (-0.450 to +0.877) | +1.03 | 0.848 | 100% | Exploratory |
| Double vs native | 694–697 | Restriction | +0.162 | +0.160 (-0.336 to +0.639) | +1.02 | 0.841 | 100% | Exploratory |
| Double vs native | 699–709 | Restriction | +0.465 | +0.466 (-0.466 to +1.355) | +1.51 | 0.916 | 100% | Supported |
| Met653 × Trp698 interaction | 685–692 | Antagonistic restriction | -0.523 | -0.531 (-1.735 to +0.669) | -2.24 | 0.115 | 100% | Exploratory |

**Note.** Positive oriented values indicate reduced RMSF (restriction). The factorial-interaction row is negative when concurrent Met653→SME antagonizes restriction relative to additivity. No isolated residue survived false-discovery-rate correction; these contiguous regions are the permitted inferential unit.

***Table S4A.*** *Cross-metric consensus regions for Cα RMSF and $S^2_{min}$.*

| Contrast | Region | Length | Direction | Mean RMSF g | Mean S²min g | Minimum RMSF P | Minimum S²min P |
|---|---|---|---|---|---|---|---|
| Met653→SME | 673–674 | 2 | Restriction | 0.92 | 0.90 | 0.834 | 0.808 |
| Trp698→KYN | 635–643 | 9 | Restriction | 1.69 | 1.71 | 0.876 | 0.820 |
| Trp698→KYN | 676–682 | 7 | Restriction | 1.58 | 1.09 | 0.836 | 0.831 |
| Double vs native | 632–633 | 2 | Restriction | 1.78 | 2.45 | 0.930 | 0.922 |
| Double vs native | 646–648 | 3 | Restriction | 1.53 | 1.26 | 0.885 | 0.806 |
| Double vs native | 657–658 | 2 | Restriction | 1.37 | 1.05 | 0.880 | 0.844 |

**Note.** Consensus required at least two adjacent residues meeting the full candidate rule in both primary profiles with concordant restriction-oriented direction.

*Table S4B. Prespecified Trp698-centered window analysis for the Trp698→KYN contrast.*

| Metric | Window | Direction | Oriented effect | Oriented g | 95% CrI | P(restriction) | Evidence |
|---|---|---|---|---|---|---|---|
| Cα RMSF | TRP698-centered window | Restriction | +0.170 | +0.94 | -0.370 to +0.706 | 0.832 | Exploratory |
| S²min | TRP698-centered window | Greater mobility / lower order | -0.042 | -0.36 | -0.386 to +0.298 | 0.341 | Uncertain |

**Note.** The window spanned residues 696–700. The two metrics did not change concordantly and therefore did not satisfy the two-metric consensus criterion.

***Table S5.*** *Condition-level projected-state occupancy, exchange, and breadth.*

| Condition | Compact (%) | Intermediate (%) | Extended (%) | Switches per 10 ns | $N_{eff}$ | Low-ΔF area ($nm^2$) |
|---|---|---|---|---|---|---|
| Native | 46.85 | 42.81 | 10.34 | 6.50 | 163.09 | 1.717 |
| Met653→SME | 70.94 | 26.84 | 2.23 | 2.37 | 116.25 | 1.602 |
| Trp698→KYN | 99.72 | 0.27 | 0.01 | 0.07 | 63.50 | 0.804 |
| Double-modified | 91.72 | 7.62 | 0.67 | 1.93 | 86.13 | 1.209 |

**Note.** Values are means across three independent trajectories. States were defined from the native oxidation-family projection and then held fixed for treatment assignment. $N_{eff}$ is the effective number of occupied bins; low-ΔF area is the area within 2RT of the projected minimum.

***Table S6.*** *Replica-level projected-state occupancy, exchange, and breadth.*

| Condition | Replica | Compact (%) | Intermediate (%) | Extended (%) | Switches per 10 ns | N_eff | Low-ΔF area ($nm^2$) |
|---|---|---|---|---|---|---|---|
| Native | 1 | 25.41 | 67.69 | 6.90 | 8.90 | 158.47 | 1.723 |
| Native | 2 | 52.13 | 38.91 | 8.96 | 6.10 | 169.00 | 1.832 |
| Native | 3 | 62.99 | 21.84 | 15.17 | 4.50 | 161.81 | 1.596 |
| Met653→SME | 1 | 18.17 | 76.98 | 4.85 | 5.10 | 181.00 | 2.612 |
| Met653→SME | 2 | 97.46 | 2.32 | 0.22 | 0.60 | 63.92 | 0.834 |
| Met653→SME | 3 | 97.18 | 1.21 | 1.61 | 1.40 | 103.83 | 1.360 |
| Trp698→KYN | 1 | 99.96 | 0.02 | 0.02 | 0.00 | 58.47 | 0.726 |
| Trp698→KYN | 2 | 99.21 | 0.78 | 0.01 | 0.20 | 77.86 | 0.961 |
| Trp698→KYN | 3 | 100.00 | 0.00 | 0.00 | 0.00 | 54.16 | 0.726 |
| Double-modified | 1 | 99.87 | 0.13 | 0.00 | 0.00 | 63.01 | 0.943 |
| Double-modified | 2 | 75.28 | 22.72 | 2.00 | 5.80 | 155.85 | 2.140 |
| Double-modified | 3 | 100.00 | 0.00 | 0.00 | 0.00 | 39.54 | 0.544 |

**Note.** The exchange-rich double-modified replica is replica 2. Replica-level values are retained to show heterogeneity and were not screened for outliers.

***Table S7.*** *Treatment-control for projected landscape properties.*

| Contrast | Metric | Estimate | 95% bootstrap CI | P(observed direction) | LOO sign stability | Evidence | Unit |
|---|---|---|---|---|---|---|---|
| Met653→SME − native | Compact occupancy | +24.09 pp | -25.06 to +63.05 pp | 0.823 | 100% | Exploratory | fraction |
| Met653→SME − native | Intermediate occupancy | -15.98 pp | -56.52 to +30.26 pp | 0.782 | 78% | Exploratory | fraction |
| Met653→SME − native | Extended occupancy | -8.12 pp | -12.88 to -3.82 pp | 1.000 | 100% | Strong | fraction |
| Met653→SME − native | Switches per 10 ns | -4.13 | -7.10 to -1.17 | 0.995 | 100% | Strong | transitions per 10 ns |
| Met653→SME − native | Low-ΔF area | -0.11 | -0.85 to +0.86 | 0.638 | 44% | Uncertain | nm² |
| Met653→SME − native | N_eff | -46.84 | -98.02 to +16.80 | 0.963 | 100% | Strong | effective bins |
| Trp698→KYN − native | Compact occupancy | +52.88 pp | +36.79 to +74.30 pp | 1.000 | 100% | Strong | fraction |
| Trp698→KYN − native | Intermediate occupancy | -42.55 pp | -67.42 to -21.63 pp | 1.000 | 100% | Strong | fraction |
| Trp698→KYN − native | Extended occupancy | -10.33 pp | -15.16 to -6.89 pp | 1.000 | 100% | Strong | fraction |
| Trp698→KYN − native | Switches per 10 ns | -6.43 | -8.83 to -4.45 | 1.000 | 100% | Strong | transitions per 10 ns |
| Trp698→KYN − native | Low-ΔF area | -0.91 | -1.07 to -0.76 | 1.000 | 100% | Strong | nm² |
| Trp698→KYN − native | N_eff | -99.59 | -111.00 to -86.34 | 1.000 | 100% | Strong | effective bins |
| Double-modified − native | Compact occupancy | +44.87 pp | +24.10 to +66.27 pp | 1.000 | 100% | Strong | fraction |
| Double-modified − native | Intermediate occupancy | -35.20 pp | -60.03 to -12.34 pp | 0.999 | 100% | Strong | fraction |
| Double-modified − native | Extended occupancy | -9.68 pp | -14.50 to -6.23 pp | 1.000 | 100% | Strong | fraction |
| Double-modified − native | Switches per 10 ns | -4.57 | -7.97 to -0.70 | 0.990 | 100% | Strong | transitions per 10 ns |
| Double-modified − native | Low-ΔF area | -0.51 | -1.15 to +0.39 | 0.889 | 100% | Exploratory | nm² |
| Double-modified − native | N_eff | -76.96 | -122.40 to -8.35 | 1.000 | 100% | Strong | effective bins |

**Note.** Simple contrasts are treatment minus native. The factorial interaction is Double − Met653→SME − Trp698→KYN + Native. Occupancy changes are percentage points (pp). The sign of compact and intermediate occupancy must be interpreted by the named state. LOO, leave one replica out.

***Table S8A.*** *Overlap and divergence of treatment and native projected distributions.*

| Treatment | Two-dimensional overlap coefficient (95% CI) | Jensen–Shannon divergence, bits (95% CI) |
|---|---|---|
| Met653→SME | 0.556 (0.252–0.647) | 0.210 (0.141–0.580) |
| Trp698→KYN | 0.195 (0.074–0.281) | 0.661 (0.552–0.859) |
| Double-modified | 0.457 (0.189–0.707) | 0.305 (0.129–0.672) |

**Note.** Overlap and Jensen–Shannon divergence were calculated on the full joint radius-of-gyration/end-to-end-distance distribution. Lower overlap and higher divergence indicate greater displacement from native.

***Table S8B.*** *Half-window projected-state occupancies for Trp698→KYN trajectories.*

| Condition | Replica | Window | Compact (%) | Intermediate (%) | Extended (%) |
|---|---|---|---|---|---|
| Trp698→KYN | 1 | Early | 100.00 | 0.00 | 0.00 |
| Trp698→KYN | 1 | Late | 99.92 | 0.04 | 0.04 |
| Trp698→KYN | 2 | Early | 98.44 | 1.54 | 0.02 |
| Trp698→KYN | 2 | Late | 99.98 | 0.02 | 0.00 |
| Trp698→KYN | 3 | Early | 100.00 | 0.00 | 0.00 |
| Trp698→KYN | 3 | Late | 100.00 | 0.00 | 0.00 |

**Note.** Early and late refer to the two halves of the 100–200 ns analysis interval. These values are sensitivity summaries; frames were not treated as independent inferential observations.
